# HOMEOSTATIC COUPLING OF CORTICAL AND BRAINSTEM DELTA RHYTHMS IN SLEEPING INFANT RATS

**DOI:** 10.64898/2026.05.11.724416

**Authors:** Midha Ahmad, Greta Sokoloff, Mark S. Blumberg

**Affiliations:** Department of Psychological & Brain Sciences, University of Iowa, Iowa City, IA 52242 USA; Iowa Neuroscience Institute, University of Iowa, Iowa City, IA 52242 USA

## Abstract

The emergence of the cortical delta rhythm (1-4 Hz) during quiet sleep (QS) is a major milestone in brain development. In rats, this milestone is achieved between 8 and 12 days of postnatal (P) age. We previously reported an age-dependent increase in PZ delta-rhythmic activity that is synchronized with cortical delta and entrained by breathing. Here, we ask whether this long-distance synchrony persists in response to perturbations to sleep homeostasis or respiration. First, using male and female P12 rats, we investigated the coupling strength between frontal cortex and PZ in response to a short but intense period of sleep deprivation. During recovery sleep, we observed a rebound in delta power in both PZ and cortex, even in the absence of increased QS duration, indicating that PZ and cortical delta power are equivalent markers of homeostatic sleep regulation. Analyses of phase-locking and lagged cross-correlation revealed persistent temporal coupling between the two rhythms such that cortical delta reliably lagged PZ delta regardless of changes in sleep pressure. Curiously, we also observed an increase in breathing depth during recovery sleep, which we confirmed in a separate cohort of pups. Next, using mild hypercapnia (5% CO_2_) to alter breathing frequency and depth, we produced decreases in cortical and PZ delta power along with decreases in the depth of breathing. These findings provide additional support for the notion that PZ and cortical delta rhythms function as distantly interconnected components within a developmentally emerging sleep-homeostatic system that is also intimately tied with the brainstem respiratory network.

**SIGNIFICANCE STATEMENT:** We reported previously that the delta rhythm that defines slow-wave sleep is not confined to the forebrain but also occurs synchronously in the medulla. This study in infant rats uses two perturbations to assess the coupling strength of cortical and brainstem delta. Using sleep deprivation and hypercapnia, we show that delta power increases or decreases in lockstep in the two regions, respectively. Our results reinforce the notion that delta across these two regions is strongly coupled and adds a new dimension to our understanding of the interconnectedness of the delta rhythm and respiration.

## INTRODUCTION

The cortical delta rhythm (1-4 Hz) is a signature feature of quiet sleep (QS) and among the earliest brain rhythms to develop (Frank & Heller, 1997; Jenni et al., 2004; Seelke & Blumberg, 2008). Nonetheless, it emerges rather late—around 2-3 months of age in humans (Jenni et al., 2004) and postnatal day 12 (P12) in rats (Gramsbergen et al., 1970; Seelke & Blumberg, 2008). In adults, delta is a key indicator of sleep pressure and depth (Borbely & Achermann, 1999; Dijk, 2009), and is thought to promote learning and neural plasticity (Girardeau & Lopes-Dos-Santos, 2021; Puentes-Mestril et al., 2019; Walker, 2009). Further, as a low-frequency rhythm, cortical delta enables long-range communication and nests higher-frequency activity (Buzsaki & Draguhn, 2004; Buzsaki & Voroslakos, 2023; Molle & Born, 2011). Given delta’s broad functional significance, its emergence is a major developmental milestone.

To begin to understand the mechanisms influencing cortical delta’s emergence in rats between P8 and P12, we recorded neural activity in the frontal cortex and parafacial zone (PZ) (Ahmad et al., 2024), a medullary region implicated in the regulation of QS in adult rodents (Anaclet et al., 2014; Anaclet & Fuller, 2017; Anaclet et al., 2012). We predicted that an age-dependent increase in PZ activity would mirror the emergence of cortical delta. But that is not what we found: Whereas PZ activity at P8 was arrhythmic during both QS and active sleep (AS), by P12 we observed periods of population-level and delta-rhythmic spiking, exclusively during QS. Even more surprising, PZ’s rhythmic spiking was phase-locked with a local delta rhythm (hereafter, PZ delta) and synchronized with cortical delta despite the long distance. Finally, PZ’s rhythmic activity increased during the post-inspiratory period, showing strong link with brainstem respiratory circuits.

Collectively, our findings suggested that PZ and cortex function together as components of a sleep-regulatory, breathing-entrained network. Accordingly, we predicted that they would exhibit similar homeostatic responses to sleep deprivation, including parallel increases in delta power during sleep rebound (Borbely & Achermann, 1999; Dijk, 2009; Tononi & Cirelli, 2006). Here, we show that PZ and cortical delta exhibit similar rebound-related increases in delta power. The rhythms remained strongly coupled, and this coupling was not affected by sleep deprivation. Moreover, cortical delta consistently lagged PZ delta, regardless of sleep pressure. We also observed, in sleep-deprived pups, unexpected increases in the depth of breathing (i.e., respiratory amplitude) that tracked the increases in delta power. The latter finding adds a new dimension to the established links between the delta rhythm and respiration in adults (Biskamp et al., 2017; Karalis & Sirota, 2022; Tort et al., 2018). We further explored these links by exposing pups to mild hypercapnia (5% CO_2_) as a means of manipulating respiration during sleep; during QS, hypercapnia reduced respiratory amplitude while also reducing delta power in PZ and cortex. Altogether, these findings indicate that at the time of delta’s developmental emergence, homeostatic regulation is coordinated by a breathing-modulated network that spans the brainstem and forebrain.

## MATERIALS AND METHODS

All procedures followed the National Institutes of Health Guide for the Care and Use of Laboratory Animals (NIH Publication No. 80–23) and were approved by the Institutional Animal Care and Use Committee at the University of Iowa.

### Subjects

Male and female Sprague-Dawley Norway rats *(Rattus norvegicus)* were used at P12–13 (hereafter, P12). Pups were housed in standard laboratory cages (48 × 20 × 26 cm) under a 12-hour light/dark cycle, with food and water provided *ad libitum*. To control for litter effects and prevent inflated statistical power (Abbey & Howard, 1973; Lazic & Essioux, 2013), littermates were always assigned to different experimental groups. Litters were culled to eight pups at P3.

PZ and cortical activity were recorded at P12 in sleep-deprived (n=8 pups, 4 males; body weight = 28.59 ± 0.58 g) and control (n=8 pups, 4 males; body weight = 29.15 ± 0.91 g) rats. In a second cohort of pups, respiration was recorded using strain-gauge plethysmography in pups that experienced the sleep-deprivation protocol (n=7 pups, 4 males; body weight = 30.08 ± 1.83 g). In a final cohort of pups, mild hypercapnia was induced to alter respiration while PZ and cortical activity were recorded (n=7 pups, 3 males; body weight = 29.97 ± 0.64 g).

### Surgery and General Experimental Procedures

As described previously (Ahmad et al., 2024), on the day of testing a pup with a visible milk band was removed from the litter and anesthetized with isoflurane gas (3-5%; Phoenix Pharmaceuticals, Burlingame, CA). Throughout the surgical procedure, the pup lay on a heated platform (Hallowell EMC, Pittsfield, MA) to maintain body temperature. The scalp was cleaned with iodine and alcohol, and a nonsteroidal anti-inflammatory drug (Carprofen, 5 mg/kg; Putney, Portland, ME) was injected subcutaneously. The scalp was removed, and a topical analgesic (0.25% bupivacaine, Pfizer, New York, NY; or lidocaine, Sparhawk Laboratories, Lenexa, KS) was applied to the skull. The skull was then dried by applying bleach with a cotton-tipped applicator. To seal the incision and prevent bleeding, Vetbond (3M, Minneapolis, MN) was applied around the perimeter of the scalp incision. Two custom-made stainless steel bipolar hook electrodes (50 μm diameter; California Fine Wire, Grover Beach, CA) were inserted into the left and right nuchal muscles and secured with collodion. The pup’s torso was wrapped in soft surgical tape (Micropore, 3M) and, to measure respiration, a piezoelectric sensor (Unimed Electrode Supplies, Surrey, UK) was secured to the back with surgical tape. In some experiments, respiration was measured instead using an elastic mercury-filled strain gauge (6.0-7.5 cm diameter; Hokanson, Inc., Bellevue, WA), stretched around the thorax and secured with soft tape. Finally, a stainless-steel head-fix apparatus (Neurotar, Helsinki, Finland) was secured to the skull using cyanoacrylate adhesive (Loctite, Henkel Corporation, Westlake, OH); the adhesive was dried rapidly using accelerant (Insta-Set, Bob Smith Industries, Atascadero, CA).

While still anesthetized, the pup was secured to a stereotaxic apparatus to complete the surgery (Stoelting Co, IL). Holes were drilled in the skull to enable subsequent electrode placement in PZ (P12: +1.80 mm rostrocaudal, relative to lambda, 1.70-1.80 mm mediolateral) and secondary motor cortex (M2; +1.0 mm rostrocaudal, relative to bregma, 1.8–2.0 mm mediolateral). A hole was also drilled over the contralateral parietal cortex for the insertion of a fine-wire thermocouple (Omega Engineering, Stamford, CT) to measure brain temperature during the acclimation period. The same hole was used for insertion of a ground/reference electrode during recording; this electrode was a chlorinated silver wire (Ag/AgCl; 0.25 mm diameter, Medwire, Mt. Vernon, NY). Mineral oil was applied to the cortical surface to prevent drying. The duration of the entire surgical procedure was approximately 30 min.

After surgery, the pup was secured to a head-fix apparatus inside a grounded Faraday cage. The pup’s torso was secured to an elevated narrow platform using surgical tape, with limbs dangling freely on each side. Heated water flowed through a radiator beneath the pup to provide warmth, and the local environment was humidified using moist sponges placed on top of the radiator. Brain temperature was monitored to ensure that it was stable at 36-37°C. The pup acclimated to the recording chamber for at least 1 h, and recording did not begin until regular sleep-wake cycles were observed. After acclimation, the thermocouple was replaced with the ground/reference electrode, at which time data acquisition began.

### Data Acquisition

Neurophysiological and respiratory data were acquired using a data acquisition system (Tucker-Davis Technologies, Alachua, FL). PZ neural data were obtained using a small-site 16-channel linear silicon electrode (A1x16-10 mm-100-177-A16; NeuroNexus, Ann Arbor, MI) inserted to a depth of 4.3-5.0 mm. Cortical LFPs were obtained using a large-site 16-channel linear silicon electrode (A1x16-10 mm-100-703-A16; NeuroNexus) inserted into M2 at a depth of 0.6–1 mm. Before insertion, electrodes were coated with fluorescent Dil (Vybrant Dil Cell-Labeling Solution, Invitrogen, Waltham, MA) for subsequent histological verification. Neural signals from the electrodes were sampled at 25 kHz using a high-pass filter (0.1 Hz) and a notch filter. Piezoelectric sensor data were sampled at 30 Hz and strain-gauge data were sampled at 1000 Hz.

#### Video recording and data synchronization

As previously described (Dooley et al., 2021; Glanz et al., 2021), body and limb movements were recorded at 100 frames/s using a digital video camera (Blackfly-S; FLIR Integrated Systems, Wilsonville, OR). Video data were acquired using SpinView software (FLIR). To synchronize video with electrophysiological recordings, an LED visible within the camera frame emitted a 100-ms pulse every 3 s. Following acquisition, a custom MATLAB script was used to insert blank frames in those rare instances when frames were dropped between LED pulses.

#### Sleep deprivation

Pups were deprived of sleep using a protocol similar to one described previously (Todd et al., 2010). Throughout the experiment, PZ and cortical activity were recorded along with respiration (using a piezoelectric sensor) and video recorded pup behavior. The experimental protocol consisted of four 30-min periods: baseline, deprivation, recovery 1 (R1), and recovery 2 (R2). The baseline period entailed uninterrupted cycling between sleep and wake. During the Deprivation period, a chilled metal spatula was gently applied to the snout around the mouth whenever cortical delta was visual observed by the experimenter, thus depriving pups of QS (but also AS as an unavoidable consequence). During R1 and R2, pups were again allowed to cycle freely between sleep and wake. Pups in the control group were left undisturbed throughout the 2-h recording period. In a follow-up experiment using a separate cohort of sleep-deprived pups, respiratory data alone were acquired using strain-gauges placed around the thorax and attached to a plethysmograph (Parks Medical Electronics Inc., Aloha, OR).

#### Hypercapnia

An additional cohort of P12 rats was exposed to mild hypercapnia (5% CO_2_ in air). For this experiment, respiration was measured using strain-gauge plethysmography and PZ and cortical activity were recorded. The recording session was divided into a 30-min baseline period of normal-air breathing, a 30-min Hypercapnia period, and a 30-min recovery period with normal-air breathing. During baseline and recovery periods, compressed air (21% O_2_, 79% N_2_) was humidified and delivered at 0.5 liters/min through a tube positioned in front of the pup’s snout. During the Hypercapnia period, everything was the same except pups breathed air supplemented with 5% CO_2_.

### Histology

At the end of recording sessions, pups were euthanized with an intraperitoneal injection ketamine/xylazine (10:1, >0.08 mg/kg). They were then perfused with 0.1 M phosphate-buffered saline, followed by 4% paraformaldehyde. Brains were extracted and then fixed in 4% paraformaldehyde for at least 24 h, after which they were transferred to a 30% sucrose phosphate-buffered solution for another 24-48 h before sectioning. Brain tissue was coronally sectioned at 80 μm using a freezing microtome (Leica Biosystems, Deer Park, IL) and mounted on gelatin-coated slides. Electrode tracks were visualized using a fluorescence microscope system (Leica Microsystems, Wetzlar, Germany). Slides were Nissl-stained with cresyl violet. Electrode placements were confirmed by overlaying the images of the fluorescent and Nissl-stained sections.

### Data Analysis

#### Pre-processing of neural data

Neural data were preprocessed using custom MATLAB scripts, as described previously (Ahmad et al., 2024; Dooley et al., 2021; Glanz et al., 2021). Raw LFP signals were down-sampled to 1000 Hz, smoothed with a 5-ms moving Gaussian kernel, and saved as binary files. Movements were detected in video recordings based on frame-by-frame changes in pixel intensity within regions of interest (ROIs; forelimbs, hindlimbs, whole-body), producing an output of real-time movement (Dooley et al., 2021; Glanz et al., 2021). Data were imported into Spike2 (Cambridge Electronic Design, Cambridge, UK) or MATLAB for further analysis.

#### Classification of behavioral state

As described previously (Ahmad et al., 2024), behavioral state was classified using a combination of respiratory and LFP activity, limb movements, and/or nuchal EMG (Dooley et al., 2021; Isler et al., 2016; Jasmeen & Pagliardini, 2017; Seelke & Blumberg, 2008),. Active wake was defined as periods when nuchal muscle tone was high and/or when multiple body parts exhibited coordinated, high-amplitude movements. As active wake transitioned to quiet wake, overt movements subsided, and muscle tone decreased. As a transition to QS occurred, respiration was regular, limb movements were absent, and PZ and cortical delta increased above the median LFP amplitude. AS began with the onset of limb twitching and included irregular respiration and arrhythmic PZ and cortical activity. If twitching ceased and at least 10 s of behavioral quiescence ensued, the AS period ended with the last twitch observed. Finally, periods of active wake, QS, and AS were required to be at least 3 s in duration to be analyzed. Microarousals (wake durations <1 s) were disregarded for the purposes of scoring changes in behavioral state.

#### Sleep pressure and rebound

Sleep pressure was operationalized as the total number of arousing cold-stimulus presentations applied during each 5-min interval of the 30-min Deprivation period. The number of presentations in each interval was averaged across pups. A repeated-measures ANOVA was used to test whether the number of applications increased over time.

To detect rebounds in sleep duration, periods of wake, QS, and AS were identified as described above. The total number of bouts for each state was calculated across the three time periods (baseline, R1, and R2) and averaged across pups; these analyses were not possible during the deprivation periods themselves due to movement artifact. To determine the total time spent in each state, the cumulative bout durations of wake, QS, and AS were calculated. Repeated-measures ANOVA was used to test for changes in duration across the three periods. The total number of bouts and the average bout duration for each state were also calculated and assessed using repeated-measures ANOVA. Log-survivor plots of bout durations, pooled across pups, were generated for wake, QS, and AS during baseline (wake: n=226 bouts; QS: n=230; AS: n=108), R1 (Wake: n=223 bouts; QS: n=193; AS: n=231), and R2 (wake: n=232 bouts; QS: n=188, AS: n=186).

To quantify rebounds in delta power, one LFP channel was selected for analysis from each PZ and cortical electrode. Channels were selected based on their anatomical location and raw delta amplitude. In the cortex, higher-amplitude delta occurred in the shallower channels. The MATLAB function, “pspectrum(),” was used to generate raw power spectra. A sampling frequency of 1000 Hz was used, and power-spectral values between 1 and 30 Hz were calculated. Power-spectral values were normalized to peak delta power, at 1.5-2.5 Hz, during baseline; normalized power was then averaged across pups. Finally, the area under the power-spectral curve within delta frequencies was calculated for each pup to compare delta power differences across the three periods. As a multiway ANOVA (3 levels within, 2 levels between) failed the Levene’s test for equality of variances, one-way repeated measures ANOVAs were instead used to test for changes in PZ and cortical delta power across time.

#### Quantifying PZ-cortical coupling for individual delta waves

In sleep-deprived adult rats, the prevalence of high-amplitude delta waves increases during recovery sleep (Vyazovskiy et al., 2007). To assess this phenomenon in sleep-deprived P12 rats, we extracted the peak amplitudes of individual delta waves in PZ and cortex using custom MATLAB scripts. To do this, LFP signals were bandpass filtered (1–4 Hz) to isolate delta waves, the peak of each wave was identified, and frequency distributions of peak amplitudes in PZ and cortex for each pup during baseline, R1, and R2 were constructed. For each pup’s frequency distribution of PZ and cortical delta during baseline, values exceeding three standard deviations above and below the mean were excluded from further analysis. From the resulting baseline distributions of delta amplitude, four percentile ranges were defined: 20^th^–40^th^, 40^th^–60^th^, 60^th^–80^th^, and ≥80^th^. These ranges served as thresholds to categorize the amplitude of individual delta waves, with waves exceeding the 80^th^ percentile designated as high amplitude. For the baseline, R1, and R2 periods, the proportion of delta peaks falling within each percentile range was computed relative to the total number of detected delta peaks.

Peri-event time histograms (PETHs) were generated to measure the degree to which high-amplitude delta peaks in PZ and cortex co-occurred during the recovery periods. For each individual cortical delta wave, the time of peak amplitude was used as the trigger, and PZ delta amplitude was plotted in relation to the trigger (window =1 s; offset =0.5 s; bin size =100 ms). The PETHs for each pup were normalized to the maximum value within the window and then averaged across pups.

To test whether the co-occurrence of high-amplitude peaks in PZ and cortex occurred above chance levels, a Monte Carlo approach was employed. Specifically, the sequence of inter-peak intervals in the cortical delta signal was shuffled 1000 times to randomize trigger times while preserving the original lengths of the intervals. (The PZ delta record was not manipulated.) Each shuffled train of cortical peaks was then jittered to remove any residual temporal alignment between signals, ensuring that any detected co-occurrence was not driven by edge effects; the value of the jitter was derived from the median inter-peak interval. The width at half-height of the average PETH for unshuffled data defined the temporal window used to calculate the probability of co-occurrence. This window was first applied to the unshuffled data to compute the observed probability, and then to each of the 1000 shuffled datasets to compute the probability expected by chance. Expected probabilities were averaged within each pup, and observed versus expected probabilities were compared across baseline, R1, and R2 periods using a 3 (period) x 2 (observed vs. expected) repeated-measures ANOVA.

#### Quantifying phase connectivity and directionality between PZ and cortical delta

Phase relations between PZ and cortical LFPs in the delta range during QS were quantified using phase differences and phase-locking values (PLVs). LFPs were first bandpass filtered in the delta range (1–4 Hz), and the Hilbert transform was applied to extract the instantaneous phase of each signal. Phase differences (cortex − PZ) were computed at each time point and summarized as normalized histograms (bin size = 12°; probabilities sum to 1) and visualized as rose plots. PLV was calculated as the absolute value of the mean phase difference (expressed as a complex vector) between the two signals. For each pup, non-uniformity of phase differences was assessed using Rayleigh’s test. Group differences in PLV between sleep-deprived (*n*=8) and control (*n*= 8) pups were evaluated using an independent-samples Mann-Whitney U test.

To obtain a measure of directionality, lagged cross-correlations were computed as previously described (Adhikari et al., 2010). Briefly, using the bandpass filtered (1–4 Hz) LFPs, instantaneous amplitude during QS was computed using the Hilbert transform. The lagged cross-correlations between PZ and cortical amplitudes were calculated using the “xcorr” MATLAB function. For each pup, the lag corresponding to the peak cross-correlation was extracted. Group-level cross-correlation values were averaged, and the distribution of peak lags was plotted across animals. Peak lags were tested against zero using a Wilcoxon matched-pairs signed-ranks test in the sleep-deprived group and a one-sample t test in controls. We then repeated this analysis in the sleep-deprived group to test if sleep pressure selectively alters directionality. For Baseline, R1 and R2 periods, peak-lags values were tested using a Friedman test.

#### Quantifying changes in depth and frequency of breathing during sleep deprivation and hypercapnia

For measures of respiratory amplitude, strain-gauge plethysmography provides a more direct measure than piezoelectric sensors. Thus, in a separate cohort of pups, we repeated the sleep-deprivation experiment using strain-gauge plethysmography. Respiratory power was calculated from the root mean square (RMS) of respiratory amplitude across sleep and wake states for the baseline, R1, and R2 periods. Changes in respiratory amplitude during R1 and R2 were assessed using a one-sample *t* test against a hypothesized value of 1 (i.e., normalized baseline) or, if normality assumptions were violated, a Wilcoxon matched-pairs signed-ranks test. This analysis was also used for the hypercapnia experiment. Additionally, respiratory frequency was quantified by detecting peaks in the respiratory rhythm and counting the total number of peaks during the baseline, hypercapnia, and recovery periods across sleep–wake states. For each period and state, the total number of peaks was divided by the total duration of that state to obtain respiratory frequency. These values were normalized to baseline, and a one-sample *t* test or Wilcoxon signed-ranks test was used as described above.

#### Quantifying changes in sleep-wake architecture during hypercapnia

Similar to the sleep-deprivation experiment, the total number of wake, QS, and AS bouts was calculated across three time periods (baseline, hypercapnia, and recovery) and averaged across pups. The average and cumulative durations of wake, QS, and AS were calculated for each time period. Repeated-measures ANOVA was used to test for changes in cumulative duration across the three periods. The total number of bouts and the average bout duration for each state were also calculated and tested using repeated-measures ANOVA.

### Statistical Analysis

All statistical analyses were conducted using SPSS 29 (IBM, Armonk, NY) or MATLAB. Statistical tests included repeated-measures ANOVA, paired *t* tests, and one-sample *t* tests. Data were tested for normality using the Shapiro-Wilks test. When normality assumptions were violated, non-parametric tests were used. When appropriate, Mauchly’s test was used to test for sphericity. In analyses where sphericity was violated, a Huynh-Feldt correction was applied to the degrees of freedom. Bonferroni corrections were used when appropriate to adjust alpha for multiple comparisons. The measure of effect size was partial eta-squared (adjusted for positive bias; (Mordkoff, 2019) for ANOVAs, Hedges’ g for *t* tests, and *r* values for Wilcoxon tests (Peres, 2025). Unless otherwise stated, means are always presented with their standard error (SEM).

Data exclusion was rare. One value that was 3 SDs above the mean was excluded as an outlier, and one pup was excluded due to persistent wakefulness.

## Results

We recorded neural activity in PZ and frontal cortex in unanesthetized, head-fixed P12 rats as they cycled between sleep and wake. Our first goal was to determine whether sleep deprivation causes parallel effects on delta power in the two structures during recovery sleep. In a separate experiment, we manipulated respiration using mild hypercapnia to further assess the strength of coupling between PZ and cortex.

### Deprived P12 rats do not exhibit sleep rebound in QS duration

In the sleep-deprived group (n=8 pups), neural activity and behavior were recorded continuously over 2 h, evenly divided into baseline, sleep deprivation, R1, and R2 periods (Fig. 1A). Sleep deprivation was achieved by applying a cold stimulus to the snout whenever the experimenter detected cortical delta. The number of stimulations required to maintain wakefulness increased significantly across the deprivation period (*F*_(5,35)_=18.04, *p*<1x10^-7^, adj. *ηp*^2^=0.68), demonstrating a monotonic accumulation of sleep pressure (Fig. 1B). The number of stimulations applied in the last 5 min of the period was twice the number during the first 5 min (*t*_(7)_=7.32, *p=.*00016, Hedges’ g=1.33).

**Figure 1.**
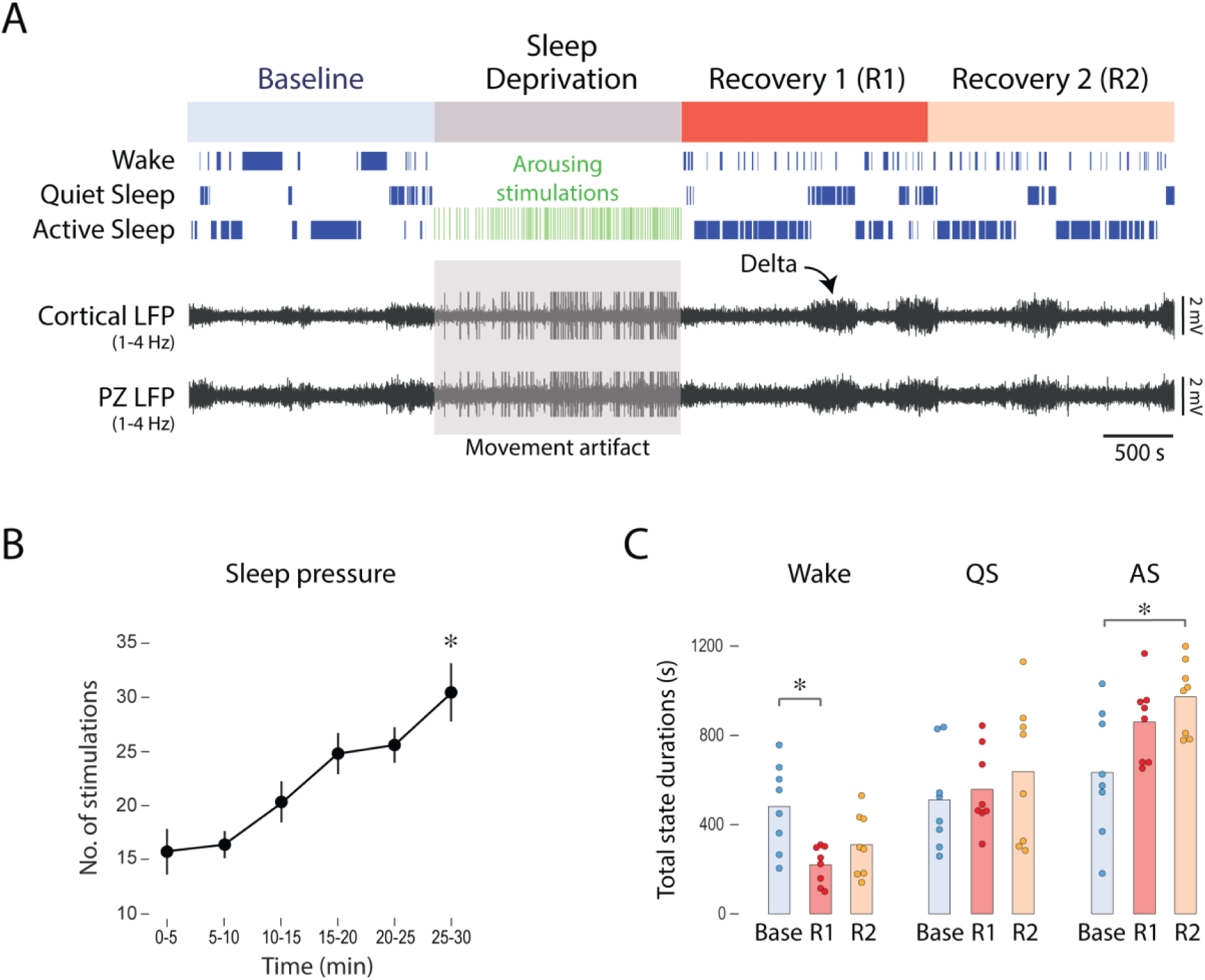
Sleep deprivation in P12 rats. ***A***, Representative data from one pup in the sleep-deprivation group. The 30-min baseline (Base), deprivation, recovery 1 (R1), and recovery 2 (R2) periods are shown; the color codes for each period are used throughout the figures. From top: Bouts of wake, quiet sleep (QS), and active sleep (AS), and cortical and parafacial zone (PZ) local field potentials (LFPs) filtered at delta frequencies (1-4 Hz). For the sleep-deprivation period, the arousing stimulations are represented as vertical green lines; the gray box indicates that the LFP data are corrupted by movement artifact. ***B***, The number of arousing stimuli presented over time during the sleep-deprivation period, indicative of accumulating sleep pressure. *n*=8 pups. The asterisk denotes significant difference from the first period, *p*<.001. ***C***, Mean total state durations for wake, QS, and AS for the baseline, R1, and R2 periods. Asterisks denote significant differences between groups, *p*s<.01.

Because the sleep cycle of P12 rats progresses sequentially from QS to AS (Seelke & Blumberg, 2008), any deprivation procedure that targets QS-related delta also necessarily deprives pups of AS. Thus, we did not expect sleep rebound to be specific to QS. Indeed, mean QS duration was unchanged during the recovery periods; instead, AS duration increased, and wake duration decreased during recovery sleep (Fig. 1C). Repeated-measures ANOVA revealed significant main effects of state (*F*_(2,14)_=14.42, *p=.*0004, adj. *ηp*^2^=0.63) and time (*F*_(2,14)=_10.48, *p=.*0017, adj. *ηp*^2^=0.54), as well as a significant state x time interaction (*F*_(4,28)_=4.43, *p=.*0067, adj. *ηp*^2^=0.30). The significant decrease in total wake duration during R1 indicates that total sleep duration increased, irrespective of the lack of selective rebounds in mean QS and AS durations during R1; indeed, there was a significant increase in total sleep duration during recovery (*F*_(2,14)=_13.86, *p=.*0005, adj. *ηp*^2^=0.62; data not shown). This result is consistent with the fact that the sleep-deprivation protocol, by targeting QS, necessarily deprives pups of both QS and AS.

Next, we assessed separately the mean number of bouts and the mean duration of each bout because those two variables, when multiplied, determine the mean total state duration. We found that the increase in total AS duration and decrease in total wake duration during recovery sleep were due primarily to an increase in the number of AS bouts and shorter mean wake durations (Fig. S1A-B). Again, QS remained unchanged across these measures.

To provide a more sensitive assessment of the statistical structure of the sleep-bout data, we constructed log-survivor plots of wake-, QS-, and AS-bout durations, pooled within baseline, R1, and R2 periods (Fig. S1). These plots show that the longest QS bouts were more consolidated during recovery sleep, thus providing evidence of an emerging selective effect of sleep deprivation on QS bout durations. In contrast, these plots show that wake-bout durations were more fragmented during recovery sleep (again indicating that total sleep-bout durations were more consolidated) while AS-bout durations were unchanged.

Recordings for the control group (n=8 pups) followed the same timeline, but the pups were undisturbed during the second 30-min period (Fig. S2A). As expected, control animals exhibited no significant changes in total state durations (*F*_(4,28)_=0.80), number of bouts (*F*_(4,28)_=2.6), or mean bout durations (*F*_(4,28)_=0.19; Fig. S2B-D).

In summary, the strongest effects of sleep deprivation on sleep architecture at P12 entailed decreases in mean wake durations during R1, indicative of increases in sleep durations. In contrast, the evidence for selective rebounds in QS and AS bouts or durations during R1 was relatively weak or absent.

### Sleep-deprived pups exhibit parallel rebounds in cortical and PZ delta power

Sleep deprivation had a substantial impact on delta rebound (Fig. 2A-B, top). Specifically, mean power spectra for cortical and PZ delta revealed ∼35% increases in power during R1, with power decreasing back toward baseline levels during R2. Mean power spectra for the control group with undisturbed sleep did not differ across periods (Fig. 2A-B, bottom).

**Figure 2.**
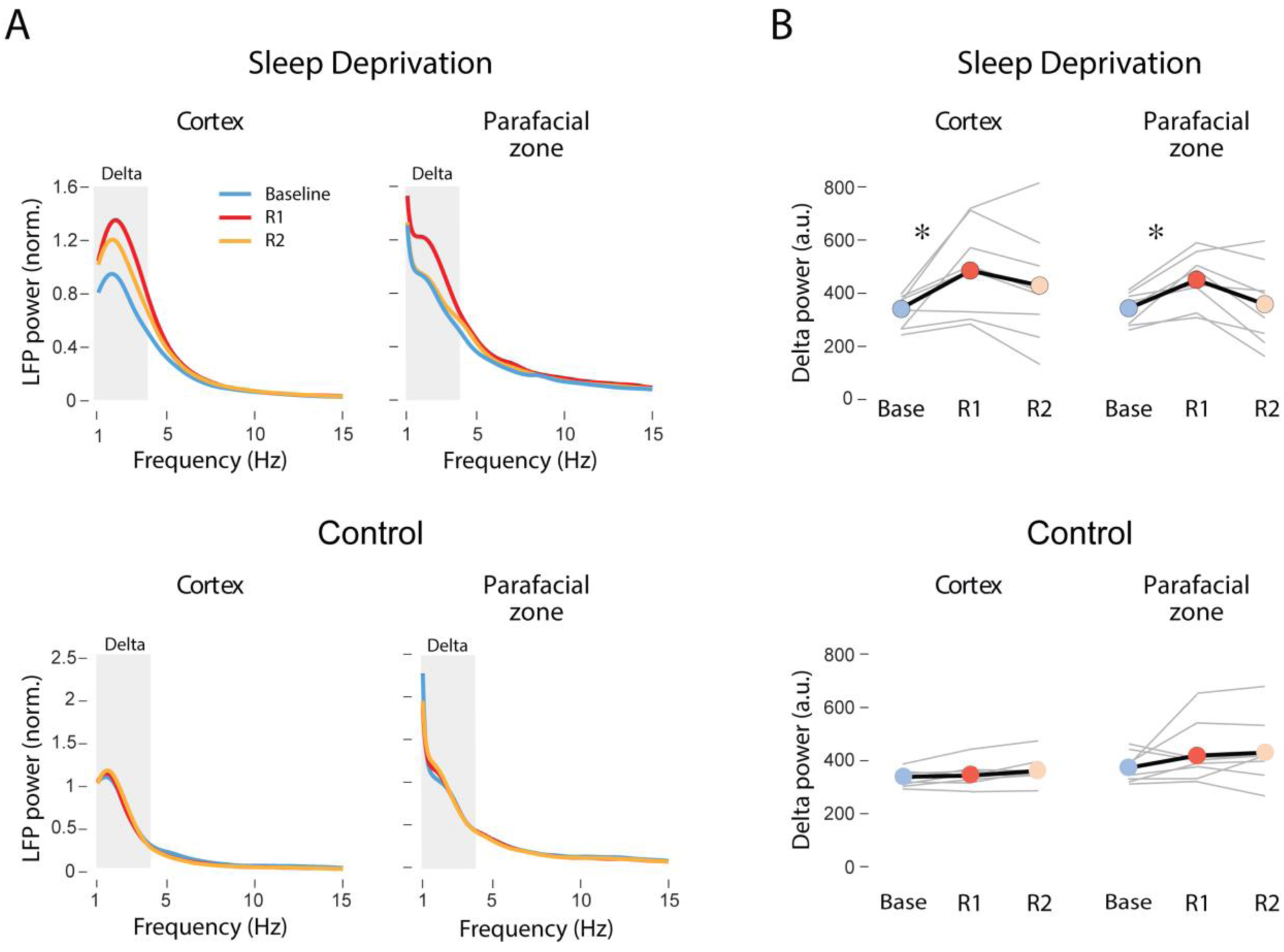
Sleep-deprived P12 rats exhibit delta-power rebound in cortex and PZ. ***A***, Mean power spectra for the frontal cortex (left) and PZ (right) during QS for baseline, R1, and R2 periods for the sleep-deprivation (top) and control (bottom) groups. Gray boxes indicate delta frequencies (1-4 Hz). Delta power was normalized to peak power (between 1.5 and 2.5 Hz) during baseline and then averaged across pups. ***B***, Mean delta power (black lines; in arbitrary units, a.u.) during QS across time for the sleep-deprivation (top) and control (bottom) groups. Gray lines show data for individual pups. Asterisks denote significant differences between time periods, *p*s<.02.

In the sleep-deprived group, there was a significant effect of time on delta power in both regions (cortex: *F*_(1.13,9.18)_=5.03, *p*=.04, adj. *ηp*^2^=0.34; PZ: *F*_(2,14)_=6.05, *p=.*013, adj. *ηp*^2^=0.39; Fig. 2B), with mean delta power during R1 being significantly greater than baseline in both cortex (t_(7)_=3.13, *p*=.016, adj. α=0.017, Hedges’ g=1.20) and PZ (t_(7)_=4.085, *p=.*005, adj. α=0.017, Hedges’ g=1.26). In contrast, during R2, mean delta power did not differ significantly from baseline in either region. In the control group, although the small changes in cortical delta power across periods were significant (*F*_(2,14)_=7.52, *p=.*006), post hoc tests were not (Fig. 2B, bottom left). There was no significant change in PZ delta power across time (*F*_(1.3,9.2)_=2.23; Fig. 2B, bottom right).

### Sleep-deprived pups exhibit increased high-amplitude delta waves in PZ and cortex during recovery sleep

In sleep-deprived adult rats, recovery sleep is characterized by the increased expression of individual high-amplitude delta waves that contribute to increases in delta power (Vyazovskiy et al., 2007). We observed a similar pattern in sleep-deprived P12 rats, with recovery sleep marked by the increased expression of high-amplitude delta waves. In both cortical and PZ LFPs, this phenomenon was expressed as parallel fluctuations in the instantaneous power envelopes (Fig. 3A).

**Figure 3.**
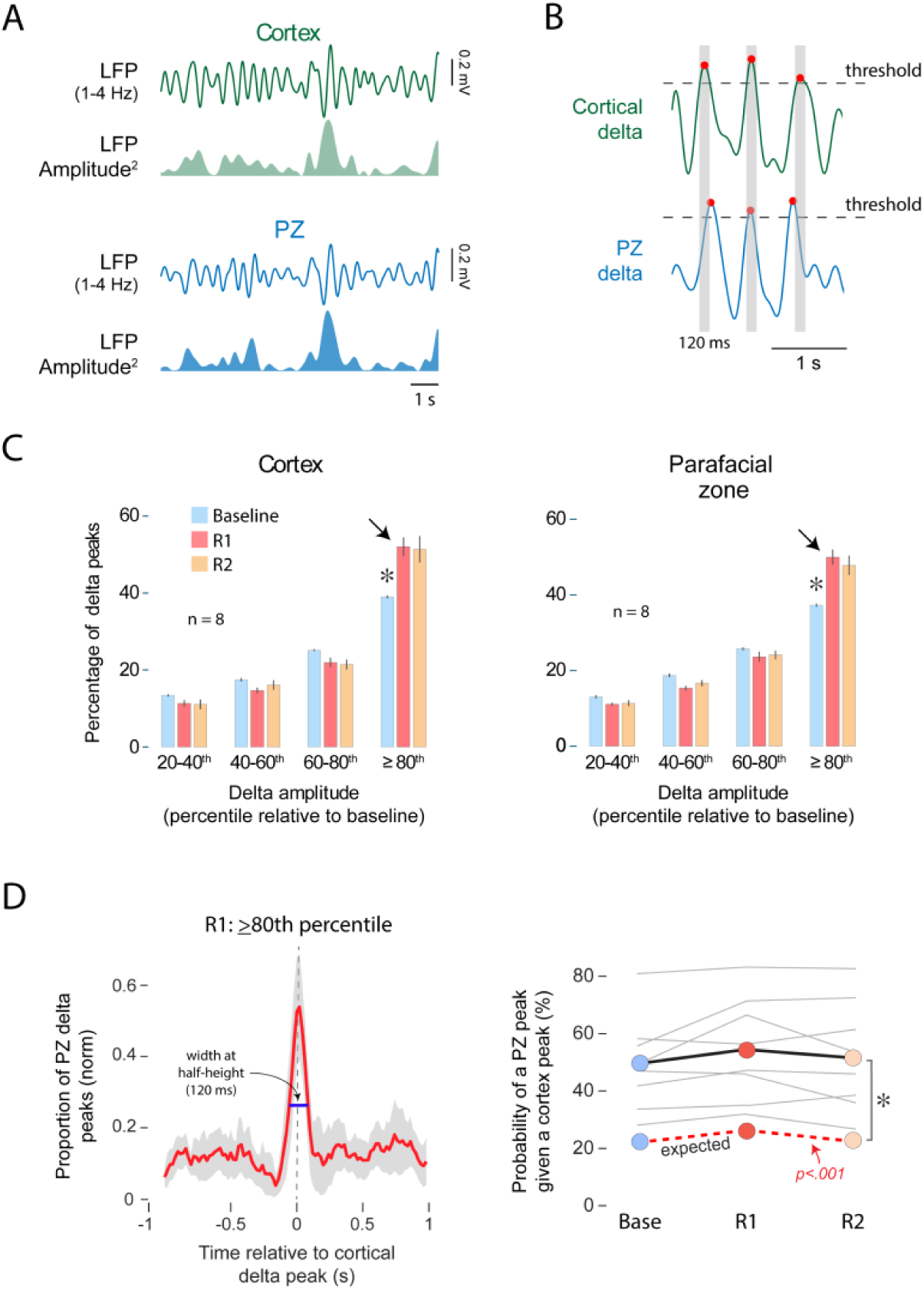
Sleep deprivation in P12 rats induces parallel increases in high-amplitude delta waves in cortex and PZ. ***A***, Representative data from one pup during QS showing instantaneous power in PZ and cortex during recovery sleep (R1). From top: Cortical LFP filtered at delta frequencies, squared instantaneous amplitude of cortical delta, PZ LFP filtered at delta frequencies, squared instantaneous amplitude of PZ delta. ***B***, Representative 2-s periods showing the co-occurrence of high amplitude delta waves in cortex (top) and PZ (bottom). Gray boxes indicate 120-ms time windows centered on peak cortical delta. ***C***, Frequency distributions of delta peaks that fell within defined threshold ranges in cortex (left) and PZ (right) for baseline, R1 and R2 periods. Asterisks denote significant difference from other two time periods, *p*s<.01. ***D***, Left: Mean peri-event histogram for PZ delta peaks triggered on cortical delta peaks. The width at half-height (120 ms) defines time windows for calculating the probability that cortical and PZ peaks co-occurred. Right: Mean observed probability (black line) of a PZ high-amplitude peak given a cortex high-amplitude. Gray lines show probabilities for individual pups. Red dotted line is the mean expected probability, calculated from shuffled data. Asterisk denotes significant difference between expected and observed probability, *p*<.01.

To quantify this effect, we first identified the peaks of individual delta waves and classified them according to their amplitudes, relative to each pup’s baseline (Fig. 3B; see Methods). This analysis confirmed a marked shift toward higher-amplitude waves during recovery sleep in both brain regions (Fig. 3C). Repeated-measures ANOVA for each brain area separately revealed significant main effects of amplitude (cortex*: F*_(3,21)_=153.93, *p* < 1x10^-7^, adj. *ηp*^2^=0.95; PZ: *F*_(3,21_=211.79, *p* < 1x10^-7^, adj. *ηp*^2^=0.96) and time (cortex: *F*_(2,14)=_169.90, *p* < 1x10^-7^, adj. *ηp*^2^=0.96; PZ: *F*_(2,14)_=63.99, *p*<1x10^-7^, adj. *ηp*^2^=0.89), as well as significant amplitude x time interactions (cortex: *F*_(6,42_=10.85, *p*<1x10^-7^, adj. *ηp*^2^=0.55; PZ: *F*_(6,42)_=13.73 *p*<1x10^-7^, adj. *ηp*^2^=0.61). Consistent with the findings in adult rats (Vyazovskiy et al., 2007), the highest-amplitude delta waves (i.e., those above the 80th percentile) were significantly more prevalent during recovery sleep in both cortex (*F*_(2,14)_=14.15, *p=.*0004, adj. *ηp*^2^=0.62) and PZ (*F*_(2,14)_=20.26, *p=.*00007, adj. *ηp*^2^=0.71).

We next determined whether the highest-amplitude waves are temporally coupled in the two brain regions. For this analysis, we constructed peri-event histograms of PZ delta peaks triggered on cortical delta peaks during R1. This analysis revealed a distinct peak at 0 s, indicating tight temporal coupling (Fig. 3D, left); approximately 60% of PZ peaks occurred within 120 ms of cortical peaks, corresponding to nearly one-quarter of a 2-Hz delta cycle. This level of co-occurrence was significantly greater than expected by chance (*F*_(1,7)_=22.32, *p=.*002, adj. *ηp*^2^=0.73; Fig. 3D, right). The same analysis during R2 yielded a similar result (data not shown).

Importantly, this coupling did not vary across time, with no significant main effect of time (*F*_(2,14)_=3.11) or a significant interaction (*F*_(2,14)_=0.78). Thus, sleep deprivation increases the *occurrence* of high-amplitude delta waves without altering coupling between the two structures.

### Cortex lags PZ in the delta-frequency range

Because high-amplitude delta waves in PZ and cortex were heterogeneous and were tightly coupled before and after sleep deprivation, we next asked whether this coupling entails a consistent phase relation between the two rhythms. First, we computed the mean phase differences between PZ and cortical delta during QS for the sleep-deprived and control groups (Fig. 4A). In both groups, phase differences were non-zero and clustered; there was a consistent phase difference (cortex – PZ) between the two rhythms of 5–30°. To assess the strength of this coupling, we computed PLVs for the two LFPs (Fig. 4B). PLVs were first tested for non-uniformity using Rayleigh’s test; all PLVs were significant (*ps*<1x10^-5^). Next, we tested for differences in PLVs between the sleep-deprived and control groups. PLVs for the sleep-deprived and control groups were not significantly different (*Z*=1.68, *p*=0.11), indicating again that the two delta rhythms are strongly coupled and that this coupling is not affected by sleep deprivation.

**Figure 4.**
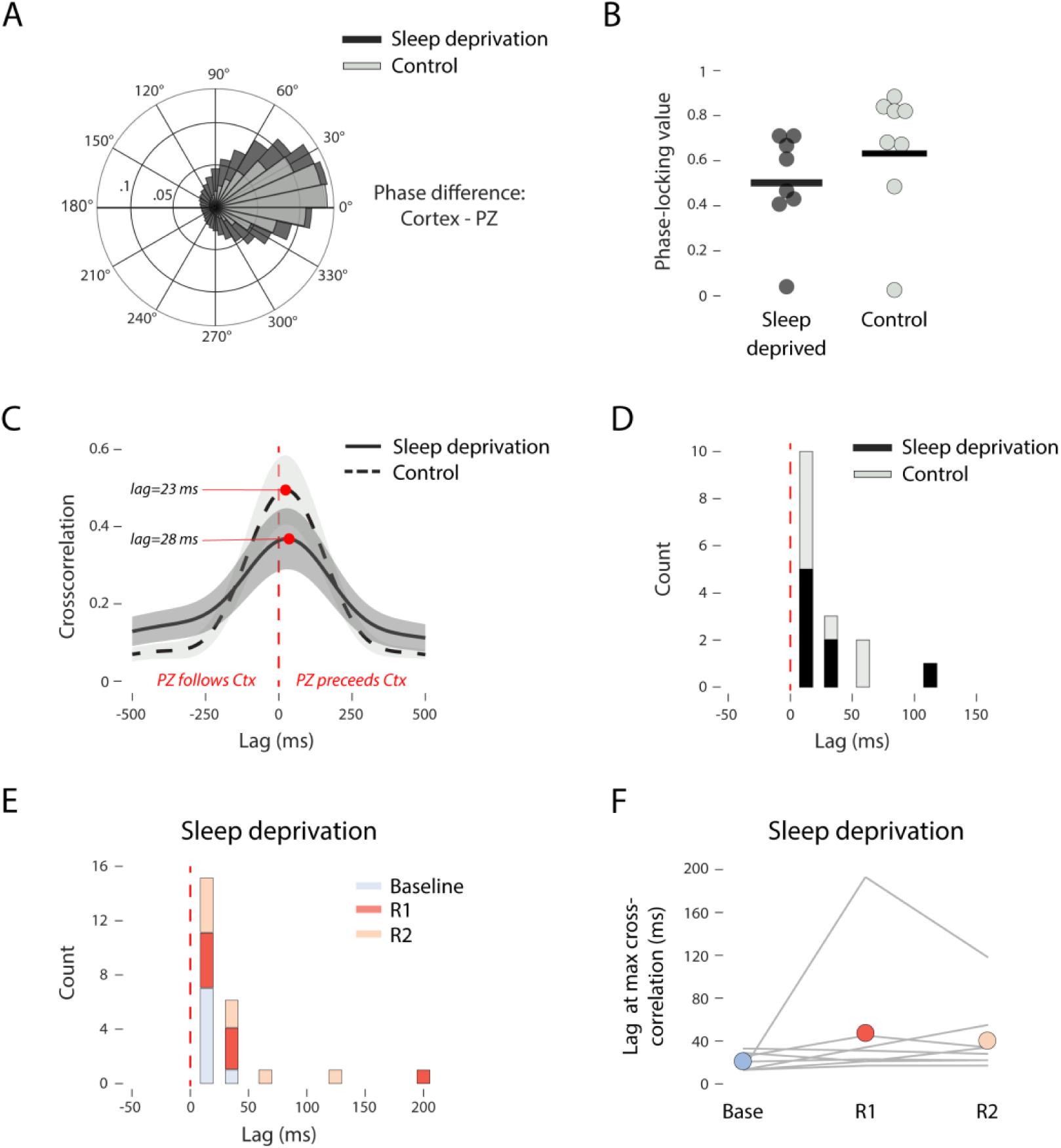
Cortical delta consistently lags PZ delta in P12 rats. ***A***, Mean phase difference between cortical and PZ delta for the sleep-deprivation and control groups. *n*=8/group. ***B***, Mean phase-locking values for cortical and PZ delta for the sleep-deprivation and control groups. Individual pups are shown, all of which exhibited significant non-uniformity (Rayleigh’s test; *p*s<1x10^-5^). ***C***, Mean lagged cross-correlations of instantaneous amplitudes for cortical and PZ delta for the sleep-deprivation and control groups. Mean lags at the maximum cross-correlations are indicated by red dots. ***D***, Count distribution of lags at maximum cross-correlation for each individual pup in the sleep-deprived and control groups. ***E***, Count distribution of lags at maximum cross-correlation for the sleep-deprivation group for the baseline, R1, and R2 periods. ***F***, Mean lags at maximum cross-correlation across time periods for the sleep-deprivation group. Gray lines represent data for individual pups.

Thus, these findings along with our previous study (Ahmad et al., 2024) provide compelling evidence of coherent PZ and cortical delta. With these results as a foundation, we next assessed the direction of information flow by computing the cross-correlations of the instantaneous amplitudes of the two delta rhythms during QS (Adhikari et al., 2010). Peak cortical-delta amplitude consistently lagged PZ-delta amplitude by an average of ∼28 ms (sleep-deprived group) and ∼23 ms (control group; Fig. 4C). Peak lags for all individual pups had positive values and were significantly different from zero for the sleep-deprived (*Z*=2.52, *p*=0.012, *r*=0.89) and control (*t*_(7)_=6.29, *p=.*003, Hedges’ g=2.26) groups (Fig 4D). Finally, we tested whether the peak lags for the sleep-deprived group varied across baseline, R1, and R2 periods: Peak lags were consistently positive at all three periods (Fig. 4E), with no significant changes over time (χ²(2)=1.87, p=0.39; Fig. 4F). Thus, cortical delta following PZ delta is a consistent feature at this age.

### Pups increase their depth of breathing during recovery sleep

In the previous experiment, it appeared that pups increased their depth of breathing during recovery sleep. Because the piezoelectric sensor used to measure breathing does not provide a direct measure of respiratory amplitude, we repeated the sleep-deprivation protocol in a new cohort of pups (n=7) but using strain-gauge plethysmography (Fig. 5). Respiratory amplitude during QS increased significantly during R1, approximately 30% over baseline values (*t*_(6)_=3.56, *p=.*012, Hedges’ g=2.06). During R2, amplitude was lower than during R1 and not significantly different from baseline (t_(6)_=1.704, *p=0*.15). Thus, increased respiratory amplitude is a reliable feature of recovery sleep at this age, mirroring the observed increase in delta power in the previous experiment.

**Figure 5.**
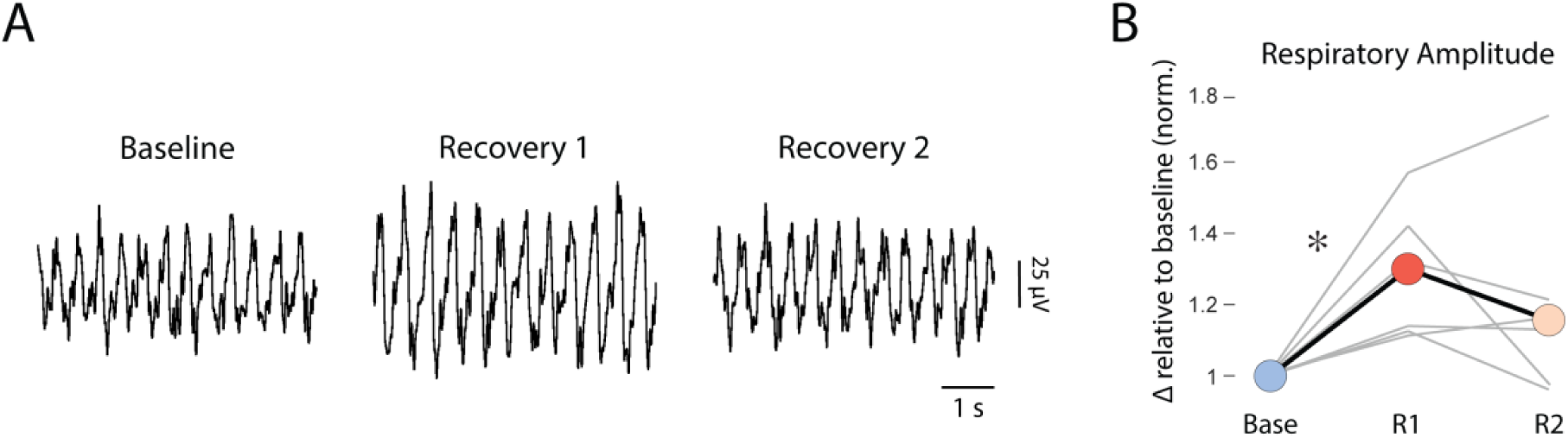
Respiratory amplitude increases during recovery sleep in P12 rats. ***A***, Representative data from one pup showing respiratory amplitude changes across time during QS. ***B***, Mean respiratory amplitude across time (black line), normalized to baseline. Gray lines represent data from individual pups. Asterisk denotes significance difference between the two time periods, *p*<.05.

### Hypercapnia suppresses PZ and cortical delta power and respiratory amplitude

As a second test of the functional coupling between PZ and cortical delta, and their relationship to respiration, we turned to a manipulation that specifically targets the respiratory system. Hypercapnia, an increase in carbon dioxide in the blood, can be produced by breathing air supplemented with CO_2_. Mild hypercapnia has variable effects on respiratory rate and amplitude in infant rats (Abu-Shaweesh et al., 1999; Putnam et al., 2005; Saetta & Mortola, 1985; Steggerda et al., 2010; Wickstrom et al., 2002), depending on such factors as age, behavioral state, duration of exposure, and anesthesia. Because sleep has not been explicitly monitored during hypercapnia in infant rats of any age, it was difficult to predict how hypercapnia would affect breathing. But given the findings above, we predicted that if hypercapnia increased or decreased breathing amplitude, we would see parallel changes in PZ and cortical delta power.

In P12 rats (n=7 pups), we assessed the effect of mild hypercapnia on respiration and PZ and cortical delta power. Pups were first allowed to breathe normal air for 30 min, followed by 30 min of breathing 5% CO_2_ and then a return to normal air for a final 30 min (Fig. 6A). Although hypercapnia produced immediate arousal, pups began cycling normally between sleep and wake within 2.5-10.8 min (mean=5.6 ± 2.8 min), including the expression of PZ and cortical delta. We quantified the effects of hypercapnia on sleep-wake architecture across the 30-min period and, despite the initial arousal, found no significant changes in total state duration (Fig. S3A; *F*_(4,24)_=2.12), number of bouts (Fig. S3B*; F*_(4,24)_=1.90), or average bout duration (Fig. S3A; *F*_(4,24)_=0.70).

**Figure 6.**
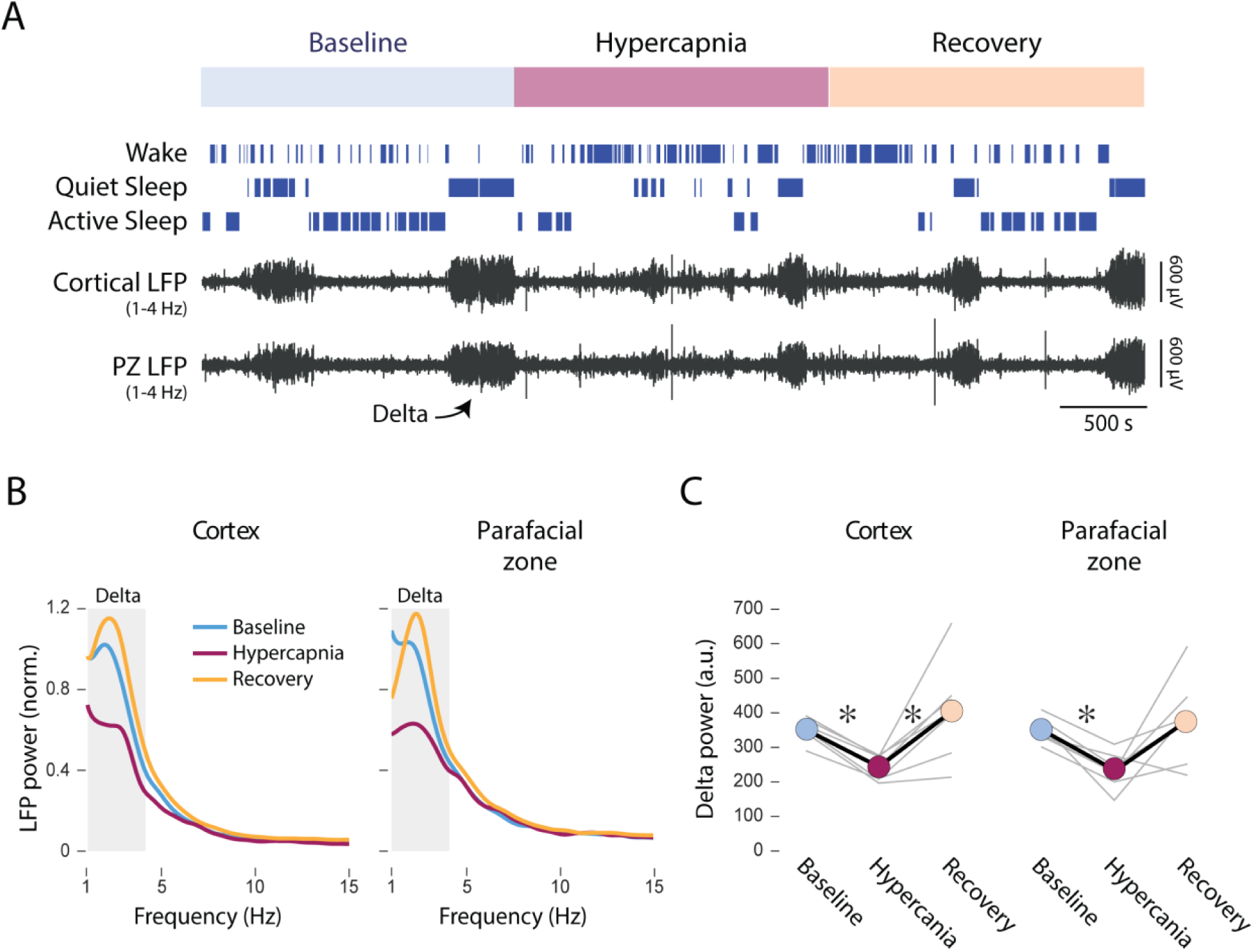
Hypercapnia suppresses QS-related cortical and PZ delta power in P12 rats. ***A***, Representative recording in a pup cycling through sleep and wakefulness. The 30-min baseline, hypercapnia (5% CO_2_), and recovery periods are shown; the color codes for each period are used throughout. From top: Bouts of wake, QS, and AS, and cortical and PZ LFPs filtered at delta frequencies (1-4 Hz). ***B***, Mean power spectra for cortex (left) and PZ (right) during baseline, hypercapnia, and recovery periods. *n=*7 pups. Gray boxes indicate delta frequencies (1-4 Hz). Delta power was normalized to peak power (between 1.5 and 2.5 Hz) during baseline, then averaged across pups. ***C***, Mean power (black lines; arbitrary units, a.u.) within delta frequencies during QS for the baseline, hypercapnia, and recovery periods. Gray lines show data for individual pups. Asterisks indicate significant differences between time periods, *p*<.01.

Even though hypercapnia did not significantly alter sleep architecture, it did affect cortical and PZ delta power during QS by ∼38% (Fig. 6B). In both regions, the decreases in delta power were reversed in the recovery period upon restoration of normal air. Statistical analysis revealed a significant effect of time on cortical (*F*_(1.08,6.48)_=9.83, *p=.*017, adj. *ηp*^2^=0.56) and PZ (*F*_(2,12)_=6.94, *p=.*01, adj. *ηp*^2^=0.43) delta power, with mean delta power during hypercapnia being significantly lower than baseline in both cortex (*t*_(6)_=12.10, *p=.*00002, adj. α=0.017, Hedges’ g=3.31) and PZ (*t*_(6)_=6.2, *p=.*0008, adj. α=0.017, Hedges’ g=2.65).

As predicted, the decreases in delta power were mirrored by a nearly 40% decrease in respiratory amplitude during the hypercapnia period (Fig. 7A). Compared to baseline, the reduction in respiratory amplitude was significant during the hypercapnia period (*t*_(6)_=3.79, *p=.*009, Hedges’ g=2.03), but not during recovery (t=1.67, *p=*0.15). The effect of hypercapnia on respiratory amplitude was specific to QS: Hypercapnia did not significantly alter respiratory amplitude during either wake or AS (*p*s<0.96; Fig. S4A). Finally, to directly compare hypercapnia-induced changes in peak PZ and cortical delta in relation to peak respiratory power, we plotted them against each other. All pups exhibited parallel decreases in delta and respiratory power in both brain structures (lower-left quadrants in Fig. 7B).

**Figure 7.**
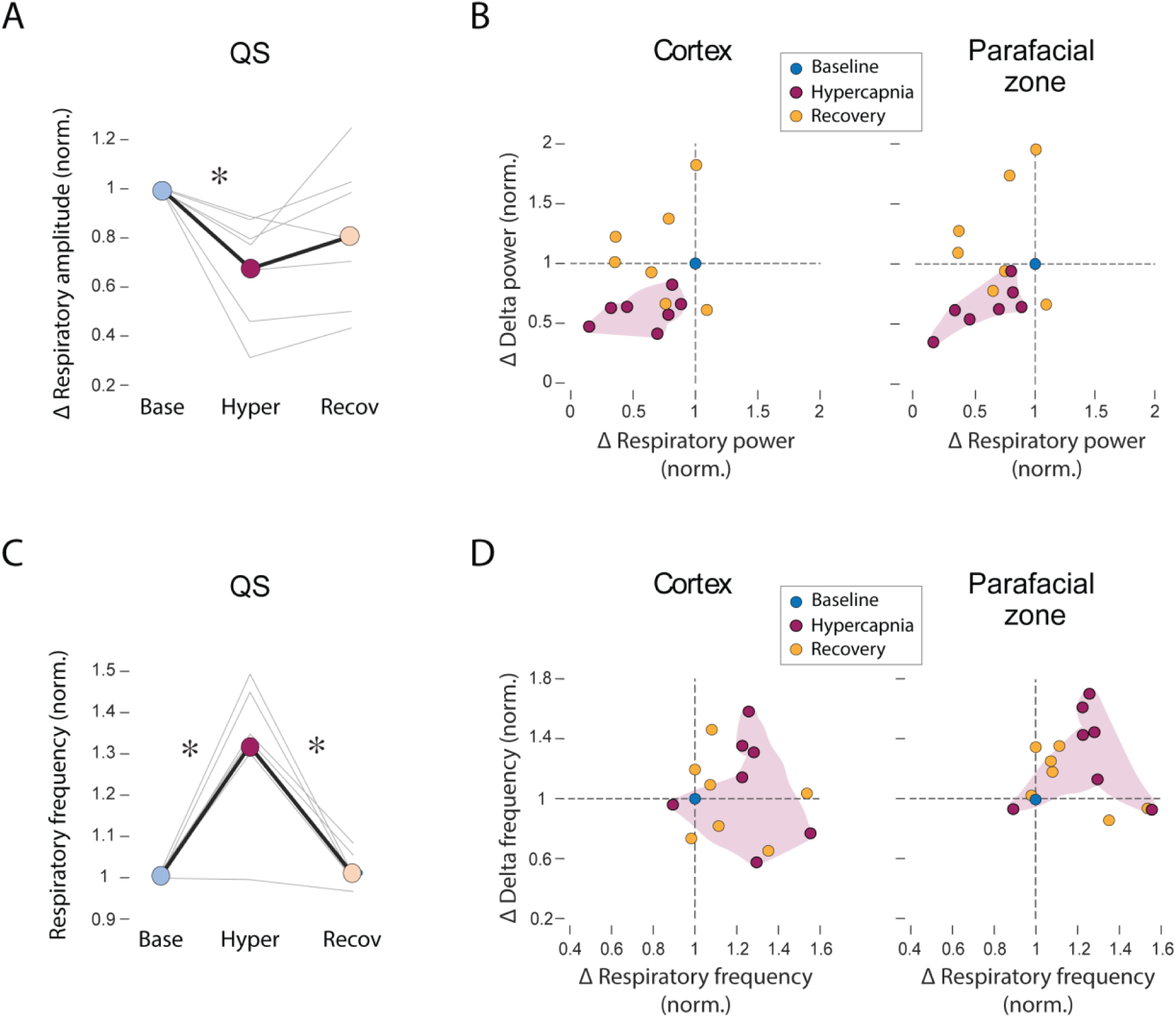
Hypercapnia evokes parallel decreases in QS-related respiratory amplitude and delta power but increases in respiratory frequency. ***A,*** Mean change in respiratory amplitude (black line) across time, normalized to baseline values. Gray lines represent changes for individual pups. *n=*7. Asterisk denotes significant difference between time periods, *p*<.01*. **B,*** Changes in peak delta power in cortex (left) and PZ (right) plotted against changes in peak respiratory power (all normalized to baseline). Each dot represents an individual pup. Purple cloud notes distribution of values during hypercapnia. ***C*** and ***D,*** Same as in ***B*** and ***C*** but for respiratory frequency.

In contrast to respiratory amplitude, hypercapnia produced, on average, ∼30-35% *increases* in respiratory frequency that were not state-dependent (Fig. 7C; Fig. S4B). The increases during QS (*t*(6)=5.28, *p=.*002, Hedges’ g=2.85), wake (*Z*=2.37; p=0.018, *r*=0.89), and AS (*t*(6)=25.23, *p*<1x10^-6^, Hedges’ g=13.48) were all significant (Fig. 7C; Fig. S4B). The decreases in respiratory frequency from hypercapnia to the recovery periods were also significant for QS (*t*(6)=5.31, *p=.*002, Hedges’ g=0.97), wake (*Z*=-2.37; p=0.018, *r*=0.89), and AS (*t*(6)=14.79, *p*=6x10^-6^, Hedges’ g=7.13). By plotting changes in PZ and cortical peak frequency against changes in respiratory frequency for each pup, we see a tendency for pups to exhibit increases in both measures (upper-right quadrant in Fig. 7D), though these patterns are not as clear as for delta and respiratory amplitude.

In summary, hypercapnia induced collective effects on PZ and cortical delta power and respiratory amplitude that were opposite to those occurring during recovery sleep after sleep deprivation. That the depressive effect of hypercapnia on respiratory amplitude occurred selectively during QS reinforces the notion that respiratory amplitude and delta power are mechanistically related.

## DISCUSSION

The present study identifies coordinated, state-dependent responses in P12 rats linking PZ and cortical delta with respiratory dynamics. First, sleep deprivation elicited parallel increases in PZ and cortical delta power during recovery sleep. Second, this rebound involved recruitment of high-amplitude delta waves in both structures, along with high-amplitude breathing. Third, PZ and cortical delta were strongly phase-locked, with cortical delta consistently lagging PZ delta. Finally, mild hypercapnia induced parallel decreases in PZ and cortical delta power along with state-dependent decreases in breathing amplitude. Thus, the synchronization of PZ and cortical delta that we reported previously at this age (Ahmad et al., 2024) persists in the face of two distinct perturbations, supporting the notion that PZ and cortical delta are coupled components of a homeostatically regulated delta-generating system. Our findings further implicate respiration as a functionally integrated element of this system at the time of its developmental onset.

### Delta-power rebound occurs as early as P12

In adult mammals, QS-related sleep homeostasis is expressed during recovery sleep as increases in delta power and/or sleep duration (Allada et al., 2017; Borbely & Achermann, 1999; Leemburg et al., 2010). In a previous study of infant rats, compensatory increases in sleep duration occurred at P12, whereas delta-power rebound was not observed until P24 (Frank et al., 1998). One explanation for the discrepancy with our findings may lie in the method used for sleep-depriving pups: Whereas the earlier study used forced locomotion or gentle handling over 3 h, we applied brief, cold stimuli to the snout, eliciting immediate and intense arousal and rapidly increasing sleep pressure over just 30 min. The efficacy of this sleep-deprivation method was first demonstrated in rats at P2 and P8, prior to the developmental emergence of cortical delta (Todd et al., 2010). Here, at P12, this method also proved effective for rapidly generating intense sleep pressure and revealing the capacity for delta homeostasis.

### Sleep deprivation does not disrupt the coupling between PZ and cortical delta

In adult rats, sleep deprivation increases the incidence of high-amplitude, synchronized slow waves during recovery sleep (Nir et al., 2011; Vyazovskiy et al., 2007). Similarly, PZ and cortical delta rebound in the present study was driven by the selective recruitment of high-amplitude delta waves. Critically, these waves were coupled in PZ and cortex, suggesting the action of a shared homeostatic network that augments delta *power* without disrupting delta *synchrony*. Supporting this interpretation, the synchronization between PZ and cortical delta remained high, with consistent, non-zero phase relationships that were unaffected by sleep deprivation. We also found that cortical delta consistently lagged PZ delta, suggesting a directional influence of the brainstem on cortical delta. Such a possibility aligns with the broader literature on a potential role for the brainstem in QS regulation (Anaclet & Fuller, 2017).

### Does breathing amplitude directly influence delta amplitude?

A surprising finding of this study is the apparent coupling of respiratory amplitude with delta power, increasing and decreasing together during recovery sleep and hypercapnia, respectively. We selected hypercapnia as a second perturbation because it provides a well-established manipulation of respiratory control (Almanza-Hurtado et al., 2022). The finding that hypercapnia produced effects opposite to those of sleep deprivation further supports a functional relationship between these signals. To the extent that hypercapnia acts primarily through respiratory pathways, this finding comes close to providing evidence of a causal effect of respiratory amplitude on delta power.

Hypercapnia also produced marked increases in respiratory frequency across all behavioral states, consistent with previous reports (Leirao et al., 2018; Molkov et al., 2014; Saetta & Mortola, 1985; Wickstrom et al., 2002). In contrast, hypercapnia’s effects on respiratory amplitude were exclusive to QS. Whereas hypercapnia typically increases the depth of breathing in adult humans (Driver et al., 2016) and rats (Leirao et al., 2018), its effect on respiratory amplitude in infant rats changes with age (Putnam et al., 2005). To our knowledge, hypercapnia’s QS-specific suppression of breathing depth has not been previously reported. Regardless, the fact that both the hypercapnia-induced changes in delta power and breathing depth are specific to QS suggests that these two processes are mechanistically linked.

That hypercapnia induces state-dependent changes in respiratory amplitude and state-independent changes respiratory frequency may reflect the separate regulatory control of these two dimensions of the respiratory system (Del Negro et al., 2018; Krohn et al., 2023). Conditions of high sleep pressure or acute hypercapnia may therefore differentially engage these systems during early development.

Together, these findings identify respiratory amplitude as a state-dependent physiological signal that is both homeostatically regulated and mechanistically coupled to delta power, revealing a previously unrecognized link between respiratory and sleep-homeostatic processes early in development.

### How is delta orchestrated across the rostrocaudal axis?

Synchronization of spatially distant PZ and cortical delta necessitates common inputs. Two possible sources of delta-driving inputs—one located rostrally and one caudally—may provide such coordination.

At the rostral end, the claustrum has emerged as a potential organizer of cortical delta (Smith et al., 2020). Across species—including rodents (Narikiyo et al., 2020), humans (Lamsam et al., 2024), and reptiles (Norimoto et al., 2020)—the claustrum is implicated in the synchronization of cortical delta. Its extensive reciprocal inhibitory cortical projections make it well-suited for this role (Jackson et al., 2020; Smith et al., 2020). An important unanswered question is whether the claustrum also coordinates delta activity beyond the cortex, including in structures like PZ.

At the caudal end, respiratory motor and sensory pathways provide a second possible source of coordination (Krohn et al., 2023; Kubin et al., 2006; Zoccal et al., 2014). There is growing evidence that respiration entrains adult brain rhythms, including cortical delta (Biskamp et al., 2017; Ito et al., 2014; Tort et al., 2018). Such entrainment may occur indirectly through olfactory pathways or directly through ascending brainstem respiratory signals (Karalis & Sirota, 2022). We showed previously that olfactory deafferentation at P12 does not disrupt cortical delta (Ahmad et al., 2024), thus supporting a direct, bottom-up effect of respiration on cortical delta.

However, respiration is unlikely to act as a stable, frequency-specific driver of delta rhythms. Whereas breathing rates vary widely across species, scaling with body mass and metabolic rate (Buzsaki et al., 2013; Fahlman et al., 2025; Glarou et al., 2025), peak delta frequency is remarkably conserved across species, occupying a narrow frequency band despite order-of-magnitude differences in body size. Respiration is therefore more likely to modulate delta dynamics than determine delta frequency itself. Moreover, given that basic aspects of state-dependent breathing already exist before P12 (Saini & Pagliardini, 2017), it is unlikely that developmental changes in respiratory function alone can explain the onset of delta.

The present finding that cortical delta consistently lagged PZ delta provides some insight into their mutual influence. However, this finding cannot be taken as evidence of PZ delta *driving* cortical delta because there may be a third region driving both at different lags (Adhikari et al., 2010). Given the short lag between these regions, and the lack of evidence of direct anatomical connectivity between them, it is possible that top-down (e.g., claustrum-mediated coordination) and bottom-up influences (e.g., respiratory entrainment) jointly synchronize PZ and cortical delta. Such bidirectional coordination could provide stability and flexibility, allowing distributed delta networks to remain synchronized while responding dynamically to physiological and homeostatic demands.

### Limitations

A limitation of the present study is its focus on a single developmental time point. Although P12 is developmentally significant, older ages should be studied to determine whether the phenomena reported here persist. A second limitation is that our assessment of PZ–cortical interactions was limited to LFP analyses because the perturbations used here—sleep deprivation and hypercapnia—introduced pronounced movement-related artifacts that prevented reliable tracking of single units across recording periods. A third limitation is that although our sleep-deprivation protocol was not intended to manipulate respiration, repeated arousing stimuli may themselves have altered breathing. Finally, stress could be a potential concern in studies involving sleep deprivation and hypercapnia; however, this concern is mitigated by the fact that rats between P4 and P14 exhibit minimal adrenocortical response to stressors (Dent et al., 2007; Walker et al., 1986).

### Conclusions and future directions

By showing that two distinct experimental perturbations were unable to dissociate the expression of PZ and cortical delta, the present findings reinforce the notion that cortical delta is closely coordinated with its brainstem counterpart. Further, evoked parallel changes in respiration and delta power adds a new dimension to the notion that breathing and delta are intimately connected.

More broadly, this study provides a foundation for understanding how sleep, respiration, brain rhythms, and long-range functional connectivity are integrated early in development. By introducing hypercapnia as a tool for manipulating respiratory–delta coupling, we establish a new approach for probing brainstem–forebrain interactions during sleep. Future work should leverage these approaches to identify the circuits and mechanisms that synchronize distributed delta-generating regions, and to determine how interactions between neural and physiological systems shape the developmental emergence of coherent brain states.

## Conflict of Interest

The authors declare no competing financial interests.

## Acknowledgments

This research was supported in part by a grant from the National Institute of Child Health and Human Development (R37-HD081168) to M.S.B.

## Notes

### Competing Interest Statement

The authors have declared no competing interest.

