## Supplemental Figures (S1-S4) for "HOMEOSTATIC COUPLING OF CORTICAL AND BRAINSTEM DELTA RHYTHMS IN SLEEPING INFANT RATS"

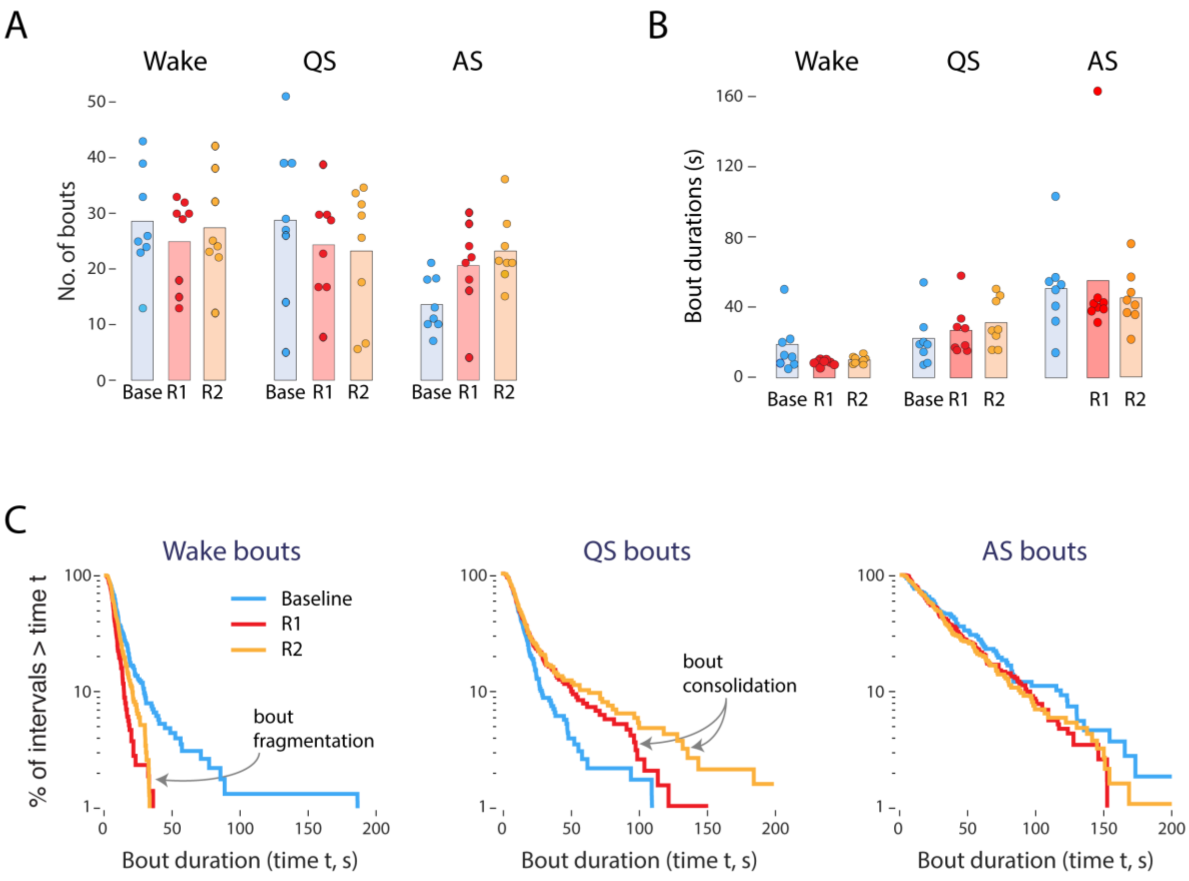


**Figure S1.** Changes in sleep-wake architecture in response to sleep deprivation in P12 rats. ***A,*** Mean number of bouts for wake, quiet sleep (QS), and active sleep (AS) across three time periods: Baseline (Base), Recovery 1 (R1), and Recovery 2 (R2). *n*=8. ***B,*** Same as in (*A*) but for mean bout durations. A significant state × time interaction was observed ($F_{(4,28)}$ = 3.94, p = .01, adj. $\eta p^2$ = 0.26), but follow-up tests were not significant ($F_{(2,14)}$ s < 3.74). ***C,*** Log-survivor plots for wake (*n*=671), QS (*n*=611), and AS (*n*=457) bouts for the baseline, R1, and R2 periods, pooled across pups.

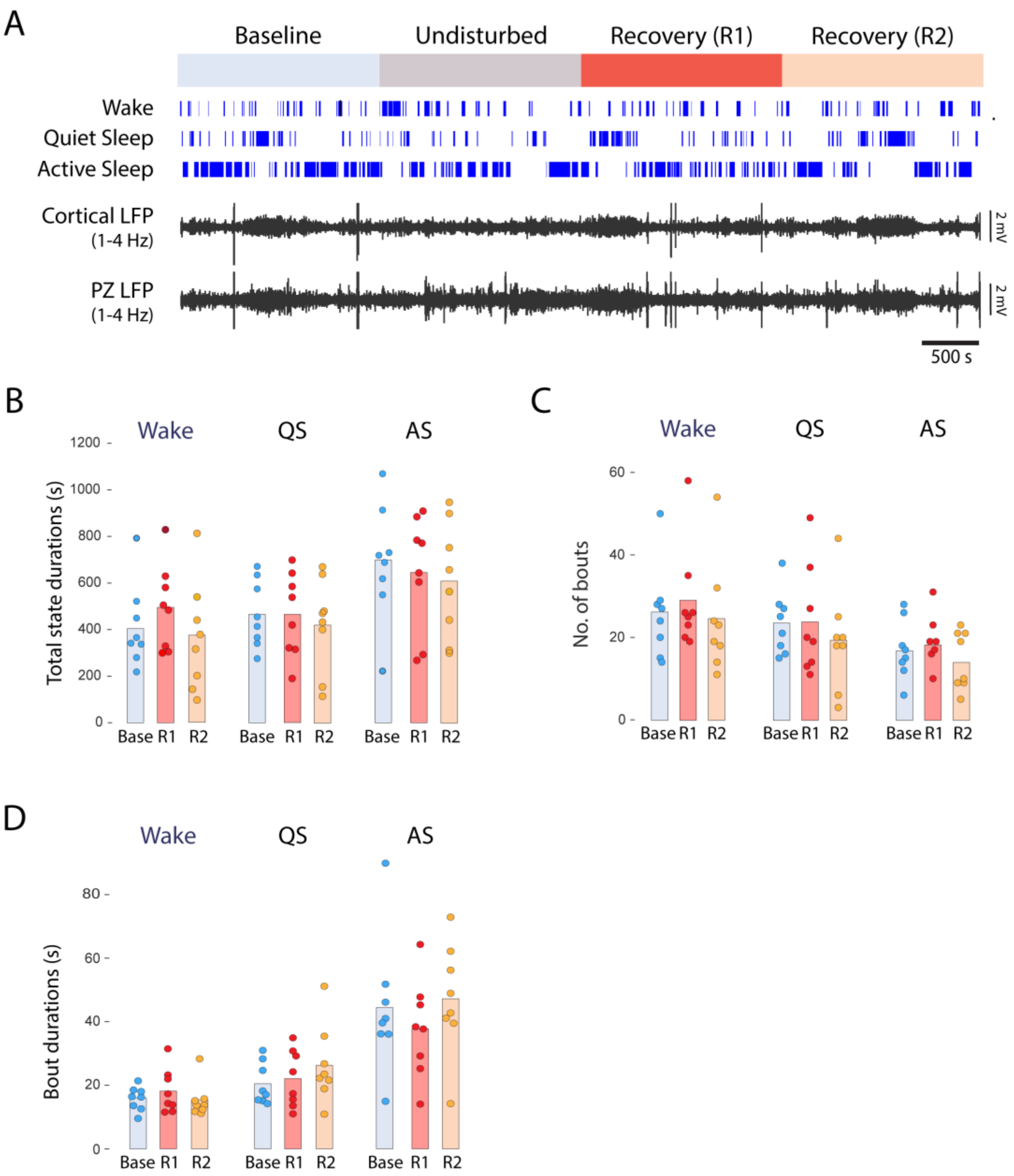


**Figure S2.** Sleep-wake architecture is unchanged in the control group with undisturbed sleep. ***A,*** Representative data from one control pup. The 30-min periods of baseline, undisturbed sleep, R1, and R2 are shown. From top: Bouts of wake, QS, and AS, cortical and PZ local field potentials (LFPs) filtered at delta frequencies (1-4 Hz). ***B,*** Mean total state durations for wake, QS, and AS for the baseline, R1, and R2 periods. *n*=8. ***C,*** Same as in (*B*) but for mean number of bouts. ***D,*** Same as in (*B*) but for mean bout durations.

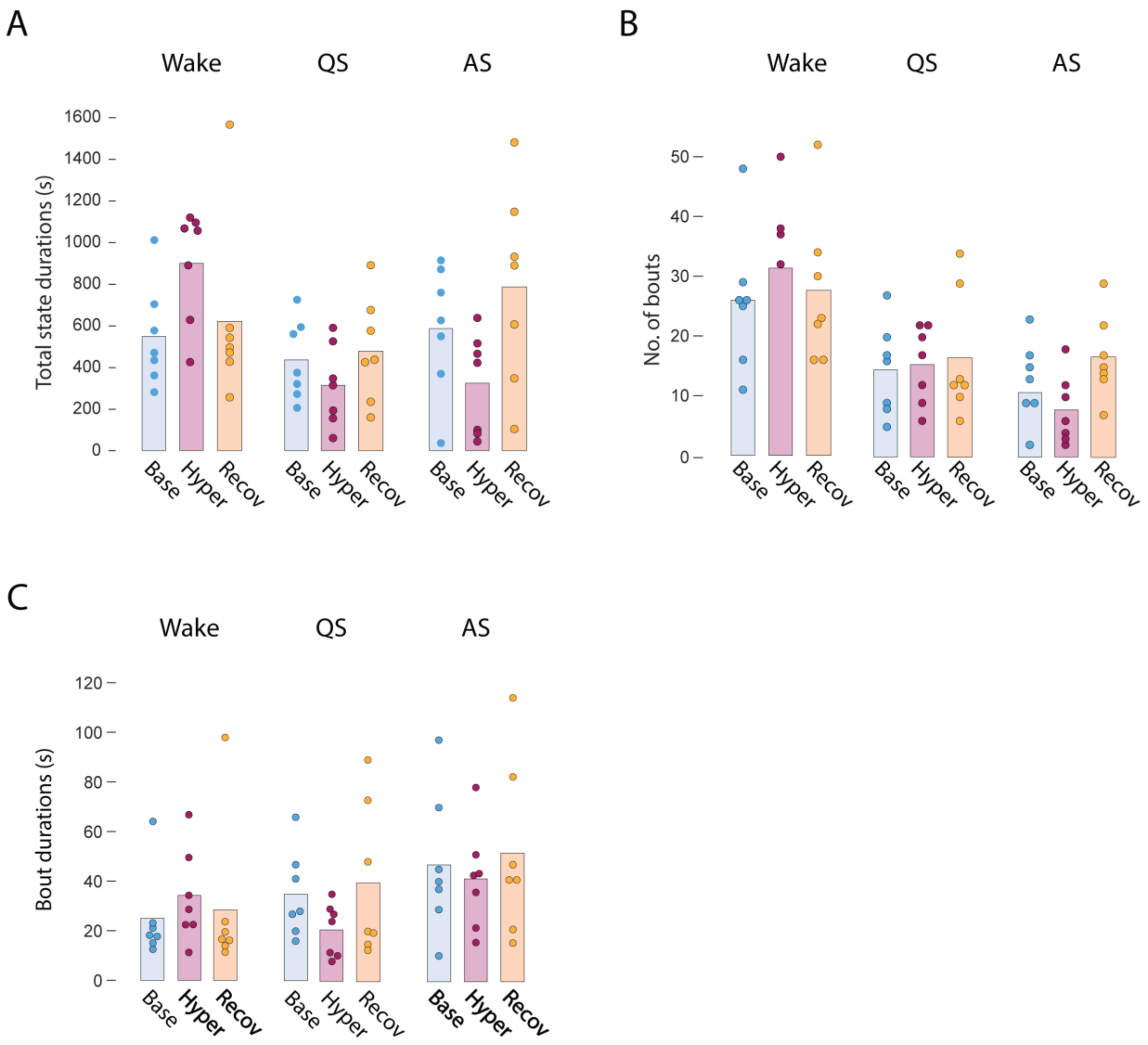


**Figure S3.** Sleep-wake architecture is unchanged in P12 rats exposed to mild hypercapnia. ***A,*** Mean total state durations for wake, QS, and AS for the baseline, hypercapnia (Hyper), and recovery (Recov) periods. *n*=7. ***B,*** Same as in (*A*) but for mean number of bouts. ***C,*** Same as in (*A*) but for mean bout durations.

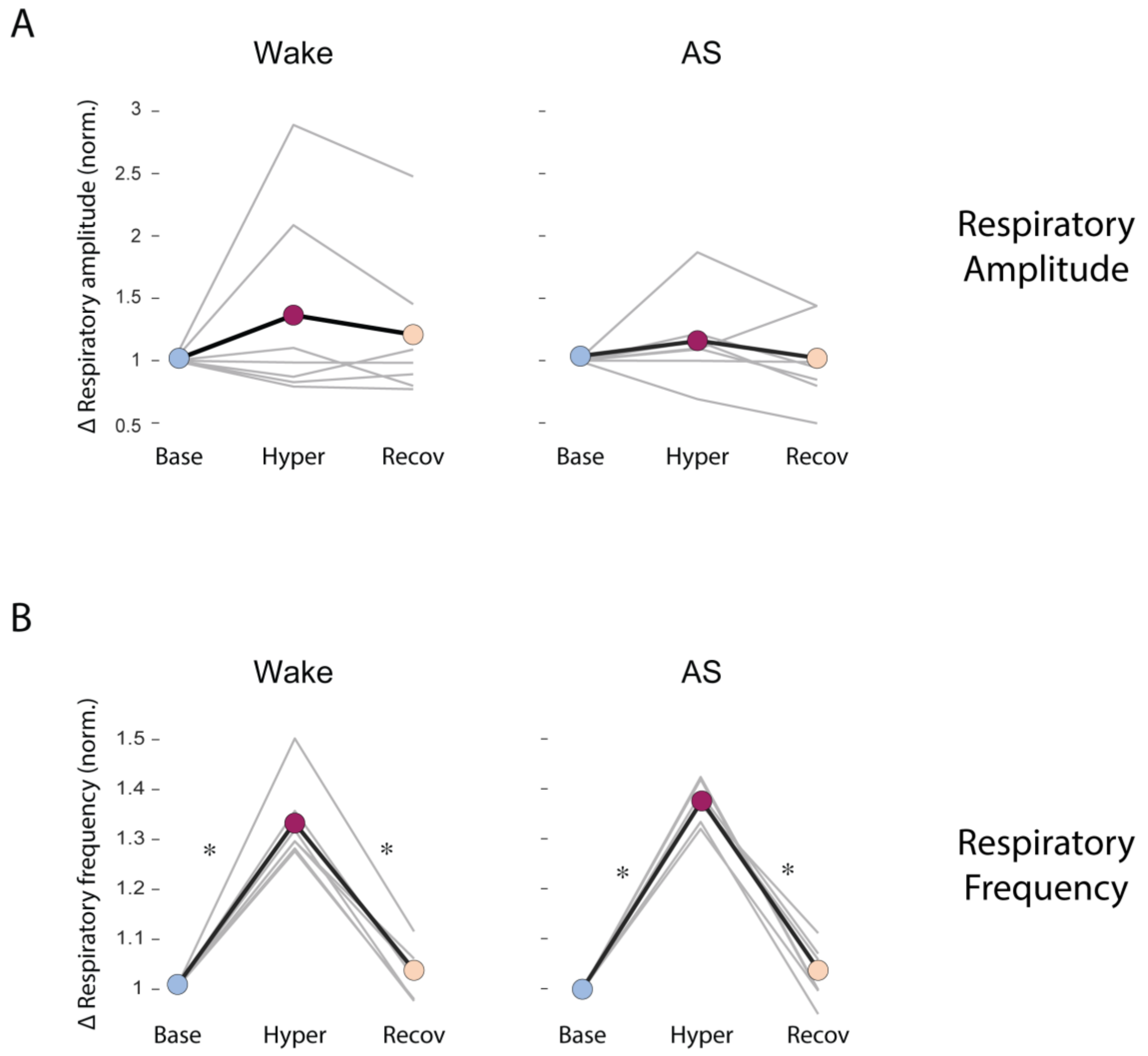


**Figure S4.** Mild hypercapnia evokes increases in respiratory frequency but not respiratory amplitude during wake and active sleep (AS). ***A,*** Mean respiratory amplitude during wake (left) and AS (right) across time, normalized to baseline. Gray lines show data for individual pups. *n*=7. ***B,*** Same as in (*A*) but for mean respiratory frequency. Asterisks denote significant differences between time periods, *ps*<.02.
